# Recent evolution of the mutation rate and spectrum in Europeans

**DOI:** 10.1101/010314

**Authors:** Kelley Harris

## Abstract

As humans dispersed out of Africa, they adapted to new environmental challenges including changes in exposure to mutagenic solar radiation. Humans in temperate latitudes have acquired light skin that is relatively transparent to ultraviolet light, and some evidence suggests that their DNA damage response pathways have also experienced local adaptation. This raises the possibility that different populations have experienced different selective pressures affecting genome integrity. Here, I present evidence that the rate of a particular mutation type has recently increased in the European population, rising in frequency by 50% during the 40,000–80,000 years since Europeans began diverging from Asians. A comparison of single nucleotide polymorphisms (SNPs) private to Africa, Asia, and Europe in the 1000 Genomes data reveals that private European variation is enriched for the transition 5’-TCC-3’→5’-TTC-3’. Although it is not clear whether UV played a causal role in the changing the European mutational spectrum, 5’-TCC-3’→5’-TTC-3’ is known to be the most common somatic mutation present in melanoma skin cancers, as well as the mutation most frequently induced *in vitro* by UV. Regardless of its causality, this change indicates that DNA replication fidelity has not remained stable even since the origin of modern humans and might have changed numerous times during our recent evolutionary history.

## Introduction

Anatomically modern humans left Africa less than 200,000 years ago and have since dispersed into every habitable environment [1]. Since different habitats have presented humans with diverse environmental challenges, many local adaptations have caused human populations to diverge phenotypically from one another. Some adaptations like light and dark skin pigmentation have been studied since the early days of evolutionary theory [2, 3, 4]. However, other putative genetic signals of local adaptation are poorly understood with respect to their phenotypic effects [5, 6, 7].

One phenotype that is notoriously hard to measure is the human germline mutation rate. It recently became possible to estimate this rate by sequencing parent-offspring trios and counting new mutations directly; however, the resulting estimates are complicated by sequencing error and differ more than 2-fold from earlier estimates inferred indirectly from the genetic divergence between humans and chimpanzees [8, 9, 10, 11]. One possible explanation for this discrepancy is a “hominoid slowdown”: a putative mutation rate decrease along the human ancestral lineage that might be related to lengthening generation time [12, 13].

A hominoid slowdown would present a caveat to the standard “molecular clock” assumption, which posits that genetic differences accumulate at a constant rate. This assumption underlies most methods for inferring demographic history from genetic variation data. There is widespread interest in using genetic variation to infer the timing of divergence and gene flow [14, 15, 16], and the accuracy of such inference is limited by the accuracy of our knowledge about mutation rates [9].

Trio-based estimates of human and chimp mutation rates have so far both fallen in the range of 1.0 *−* 1.25 *×* 10^−8^ mutations per site per generation [10, 11, 17]. However, the two species appear to differ in the distribution of *de novo* mutations between the male and female germlines and among different mutation types (e.g. a higher proportion of chimp mutations are CpG transitions) [17]. These patterns suggest that there has been some degree of mutation rate evolution since the two species diverged.

To my knowledge, previous studies have presented no evidence of mutation rate evolution on a timescale as recent as the human migration out of Africa. Most human trios that have been sequenced are European in origin (see Supporting Information S9), meaning that there exist few measurements of *de novo* mutation patterns on diverse genetic backgrounds. However, there is some reason to suspect that mutation rates might have changed due to recent regional adaptations affecting DNA repair. SNPs that affect gene expression in DNA damage response pathways show evidence of recent diversifying selection, exhibiting geographic frequency gradients that appear to be correlated with environmental UV exposure [7]. I sought to test whether mutation rates vary between populations using rare segregating SNPs that arose as new mutations relatively recently [11, 18], examining the 1000 Genomes data for mutation spectrum asymmetries that could be informative about human mutation rate evolution.

## Results

### Mutation spectra of continent-private variation

To test for differences in the spectrum of mutagenesis between populations, I compiled sets of population-private variants from the 1000 Genomes Phase I panel of 1,092 human genome sequences [18]. Excluding singletons and SNPs with imputation quality lower than RSQ = 0.95, which might be misleadingly classified as population-private due to imputation error, there remain 462,876 private European SNPs (PE) that are variable in Europe but fixed ancestral in all non-admixed Asian and African populations, as well as 265,988 private Asian SNPs (PAs) that are variable in Asia but fixed ancestral in Africa and Europe. These SNPs should be enriched for young mutations that arose after humans had already left Africa and begun adapting to temperate latitudes. I compared PE and PAs to the set of 3,357,498 private African SNPs (PAf) that are variable in the Yorubans (YRI) and/or Luhya (LWK) but fixed ancestral in Europe and Asia. One notable feature of PE is the percentage of SNPs that are C*→*T transitions, which is higher (41.01%) than the corresponding percentages in PAs (38.99%) and PAf (38.29%).

Excess C*→*T transitions are characteristic of several different mutagenic processes including UV damage and cytosine deamination [19]. To some extent, these processes can be distinguished by partitioning SNPs into 192 different context-dependent classes, looking at the reference base pairs immediately upstream and downstream of the variable site [20]. For each mutation type *m* = *B*_5′_*B_A_B*_3′_ → *B*_5′_*B_D_B*_3′_ and each private SNP set *P*, I obtained the count *C*_*P*_ (*m*) of type-*m* mutations in set *P* and used a *χ*^2^ test to compare *C*_PE_(*m*) and *C*_PAs_(*m*) to *C*_PAf_(*m*).

As shown in Figure 1A, the strongest candidate for mutation rate change is the transition 5’-TCC-3’*→*5’-TTC-3’ (hereafter abbreviated as TCC*→*T). Combined with its reverse strand complement 5’-GGA-3’*→*5’-GAA-3’, TCC*→*T has frequency 3.32% in PE compared with 1.98% in PAf and 2.04% in PAs. Several other C*→*T transitions are also moderately more abundant in PE than PAf, in most cases flanked by either a 5’ T or a 3’C.

**Fig. 1.**
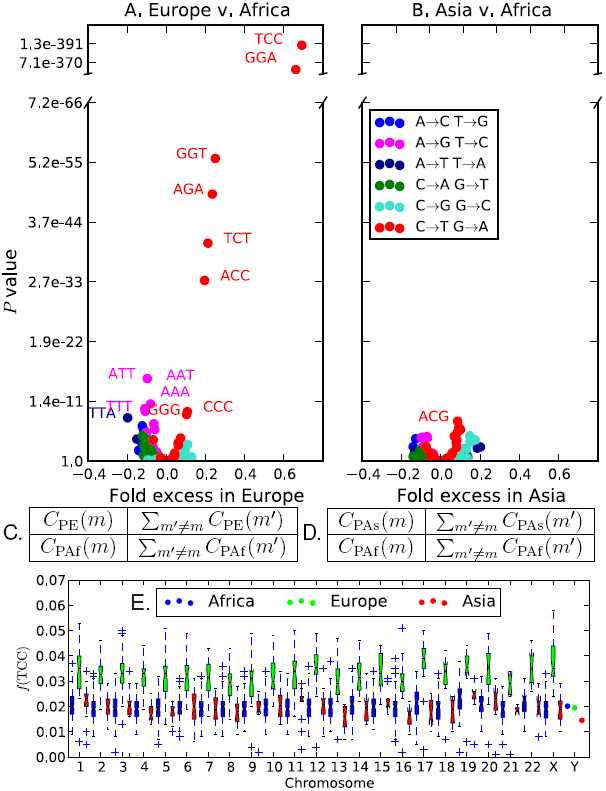
Overrepresentation of 5’-TCC-3’*→*5’-TTC-3’ within Europe. Panels A,B: The *x* coordinate of each point in gives the fold frequency difference (*f*_PE_(*m*) *– f*_PAf_(*m*))*/f*_PAf_(*m*) (resp. (*f*_PAs_(*m*) – *f*_PAf_(*m*))*/f*_PAf_(*m*)), while the *y* coordinate gives the Pearson’s *χ*^2^ *p*-value of its significance. Outlier points are labeled with the ancestral state of the mutant nucleotide flanked by two neighboring bases, and the color of the point specifies the ancestral and derived alleles of the mutant site. Panels C and D show the *χ*^2^ contingency tables used to compute the respective *p* values in Panels A and B. Panel E shows the distribution of *f* (TCC) across bins of 1000 consecutive population-private SNPs. Only chromosome-wide frequencies are shown for Chromosome Y because of its low SNP count.

The TCC*→*T frequency difference holds genome-wide, evident on every chromosome except for chromosome Y, which has too little population-private variation to yield accurate measurements of context-dependent SNP frequencies (Figure 1E). The most parsimonious explanation is that Europeans experienced a genetic change increasing the rate of TCC*→*T mutations. C*→*T transitions may not be the only mutations that experienced recent rate change; for example, TTA*→*TAA mutations appear to be less abundant in Europe than in Africa. Several CpG transitions including ACG*→*ATG appear to have higher frequencies in Asia than Africa, but these differences disappear when the spectrum is folded to e.g. identify ACG*→*ATG with the inverse mutation ATG*→*ACG (Supporting Information Section S5). This is not surprising given that high CpG mutation rates makes these sites especially susceptible to ancestral state miscalls [21].

If mutation type *m* occurs at a higher rate in Europe than in Asia, a European haplotype should contain excess type-*m* derived alleles compared to an Asian haplotype. This prediction is tested in Section S1 of the Supporting Information. The results suggest that many mutation types occur at slightly higher rates in Europe compared to Asia, with C*→*T transitions, particularly TCC*→*T, showing the strongest signal of rate differentiation. This asymmetry cannot be explained by a demographic event such as a population bottleneck; however, it should be interpreted with caution because many bioinformatic biases have the potential to confound this test. Prüfer, et al. used a similar technique to quantify divergence between archaic and modern genomes and found that branch length differences between sequencing batches often exceeded branch length differences between populations [22]. In addition, because the 1000 Genomes Phase I dataset is heavily imputed and contains more European genomes than Asian genomes, rare European variants might be ascertained more completely than rare Asian variants. This could produce a false overall excess of European derived alleles, but seems unlikely to elevate the discovery rate of TCC*→*T in Europe relative to other mutation types.

### Robustness to sources of bioinformatic error

Figures 1, S2, and S3 suggest that the human mutation rate is remarkably labile, with significant change having occurred since the relatively recent European/Asian divergence. In this section, I summarize evidence that this conclusion is not founded on bioinformatic artifacts. I focus on confirming the veracity of the TCC*→*T excess in Europe, but do not discount the possibility that other mutation types might have experienced smaller rate changes.

To rule out the possibility that TCC*→*T excess in Europe is a bioinformatic artifact specific to the 1000 Genomes data, I reproduced Figure 1A,B in a set of human genomes sequenced at high coverage using Complete Genomics technology (Supporting Information Section S3) [23]. I also folded the context-dependent mutation frequency spectrum to check for effects of ancestral misidentification (Supporting Information Section S5). Finally, I partitioned the 1000 Genomes data into bins based on GC content, sequencing depth, and imputation accuracy, finding that the TCC*→*T excess in Europe was easily discernible within each bin (Supporting Information Sections S6, S7, and S10). Three other C*→*T transitions (TCT*→*TTT, ACC*→*ATC, and CCC*→*CTC) are also more abundant in Europe than Africa across a broad range of GC contents and sequencing depths. In contrast, genomic regions that differ in GC content and/or sequencing depth show little consistency as to which mutation types show the most frequency differentiation between Africa and Asia.

As mentioned previously, singleton variants (minor allele count = 1) were excluded from all analyses. When singletons are included, they create spurious between-population differences that are not reproducible with non-singleton SNPs (Supporting Information Section S8). This is true of both the low coverage 1000 Genomes dataset and the smaller, higher coverage Complete Genomics dataset, suggesting that singletons are error-prone even in high coverage genomes.

A particularly interesting class of singletons are *de novo* mutation calls in trios. Barring bioinformatic problems, counting these mutations should yield an accurate estimate of the current human mutation rate and spectrum. I compared PE and PAs to the *de novo* mutation calls from 82 Icelandic trios by rank-ordering mutation types in each dataset from most frequent to least frequent (Supporting Information Section S9) [10]. In PE, TCC*→*TTC and its complement GGA*→*GAA are the 16th and 17th most common SNP types, respectively, whereas they are only ranked 27th and 32nd in PAs. In the Icelandic trios, TCC*→*TTC and GGA*→*GAA are ranked 20th and 18th. By this measure, the Icelandic mutations resemble PE more closely than PAs, as expected for European trios.

### Antiquity of the European mutation rate change

The 1000 Genomes Phase I dataset contains samples from five European sub-populations: Italians (TSI), Spanish (IBS), Utah residents of European descent (CEU), British (GBR), and Finnish (FIN). All of these populations have elevated TCC*→*T frequencies, suggesting that the European mutation rate changed before subpopulations diversified across the continent. To assess this, I let P_total_ denote the combined set of private variants from PE, PAs, and PAf, and for each haplotype *h* let P_total_(*h*) denote the subset of Ptotal whose derived alleles are found on haplotype *h*. *f*_*h*_(TCC) then denotes the frequency of TCC*→*T within P_total_(*h*). For each 1000 Genomes population *P*, Figure 2 shows the distribution of *f*_*h*_(TCC) across all haplotypes *h* sampled from *P*, and it can be seen that the distribution of *f* (TCC) values found in Europe does not overlap with the distributions from Asia and Africa. In contrast, the four admixed populations ASW (African Americans), MXL (Mexicans), PUR (Puerto Ricans), and CLM (Colombians) display broader ranges of *f* (TCC) with extremes overlapping both the European and non-European distributions. The African American *f* (TCC) values are only slightly higher on average than the non-admixed African values, but a few African American individuals have much higher *f* (TCC) values in the middle of the admixed American range, presumably because they have more European ancestry than the other African Americans who were sampled.

**Fig. 2.**
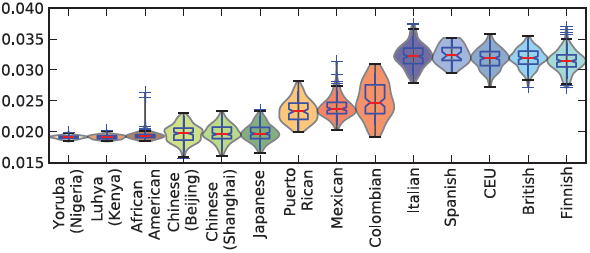
Variation of *f* (TCC) within and between populations. This plot shows the distribution of *f* (TCC) within each 1000 Genomes population, i.e. the proportion of all derived variants from PA, PE, and PAf present in a particular genome that are TCC → T mutations. There is a clear division between the low *f* (TCC) values of African and Asian genomes and the high *f* (TCC) values of European genomes. The slightly admixed African Americans and more strongly admixed Latin American populations have intermediate *f* (TCC) values reflecting partial European ancestry.

Within Europe, Figure 2 shows a slight *f* (TCC) gradient running from North to South; the median *f* (TCC) is lowest in the Finns and highest in the Spanish and Italians. In this way, TCC*→*TTC frequency appears to correlate negatively with recent Asian co-ancestry (Supporting Information Section S2).

To roughly estimate the time when the TCC*→*T rate increased, I downloaded allele age estimates that were generated from the Complete Genomics data using the program ARGweaver (http://compgen.bscb.cornell.edu/ARGweaver/CG_results/) [24]. Based on these estimates, TCC*→*T rate acceleration appears to have occurred between 25,000 and 60,000 years ago, not long after Europeans diverged from Asians (Supporting Information Section S4). In the 1000 Genomes, data, TCC*→*T frequency differentiation is greatest for private alleles of frequency less than 0.02 (Supplementary Figure S6B).

It is hard to tell from current data whether skin lightening predated TCC*→*T acceleration in Europe. A 7,000-year-old Early European farmer was found to be homozygous for the skin-lightening SLC24A5 allele [25], suggesting that light skin was relatively prevalent by 7,000 years ago and could have originated much earlier. An attempt to date the origin of this allele yielded a 95% confidence interval of 6,000 to 38,000 years ago [26], which overlaps with the time interval when the TCC*→*T rate appears to have accelerated.

### Reversal of TCC*→*T transcription strand bias in Europe

Transcribed genomic regions are subject to transcription-coupled repair (TCR) of damaged nucleotides that occur on the sense DNA strand. This can lead to patterns of strand bias that contain information about underlying mutational mechanisms. For example, CpG transitions generally result from deamination damage to the cytosine rather than the guanine, and damaged Cs that occur on the transcribed strand are repaired more often than damaged Cs occurring on the nontranscribed strand. As a result, CpG transitions in genic regions are usually oriented with the C*→*T change on the nontranscribed strand [27].

To assess the strand bias of genic TCC*→*T mutations and look for strand bias differences between populations, I counted the occurrences of each A/C ancestral mutation *m* from each private SNP set *P* on transcribed gene strands versus nontranscribed gene strands, denoting these counts **T**(*P, m*) and **N**(*P, m*), respectively. Strand biases **S**(*P, m*) = **N**(*P, m*)*/***T**(*P, m*) were compared between populations using a *χ*^2^ test (Figure 3). Private Asian and African TCC*→*T SNPs were found exhibit the strand bias that is typical of A/C*→*G/T mutations [28], with the C*→*T change usually affecting the antisense strand and the G*→*A change usually affecting the sense strand. In contrast, private European TCC*→*T SNPs exhibit no discernible strand bias; the C*→*T change affects the sense strand about 50% of the time (Figure 3E). TCC*→*T is the only mutation type that exhibits a significant strand bias difference between populations at the level *p <* 0.01.

**Fig. 3.**
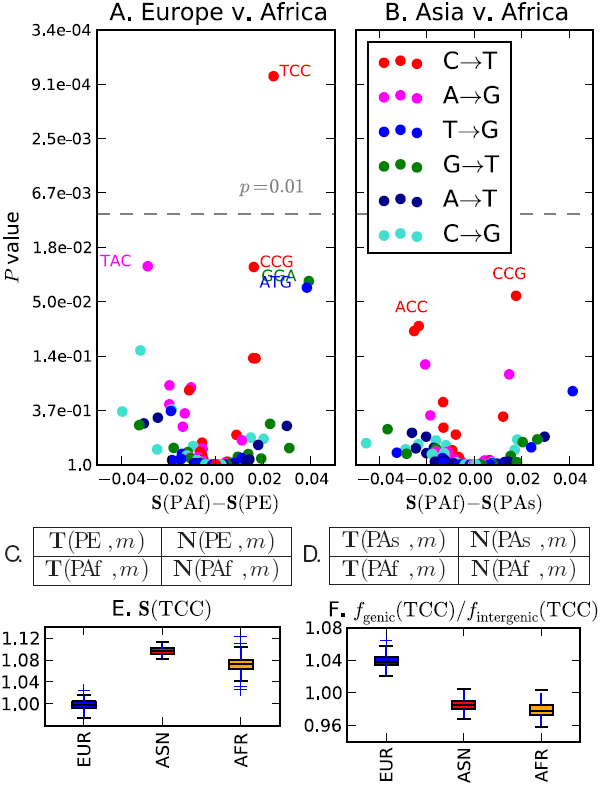
Differences in transcriptional strand bias. Each point in panels A and B represents a mutation type with an A or C ancestral allele. The *x* coordinate of each point in panel A is the PAf strand bias minus the PE strand bias; similarly, the *x* coordinates in panel B describe the PAf strand bias minus the PAs strand bias. The *y* coordinate of each point is the *χ*^2^ *p*-value of the strand bias difference. At the *p* = 0.01 significance level (grey dashed line), only TCC → T has a significant strand bias difference between Europe and Africa, while no mutation type significantly differs in strand bias between Asia and Africa. Panel C shows the variance of strand bias in each population across 100 bootstrap replicates. Similarly, Panel D shows the distribution across bootstrap replicates of the ratio between genic *f* (TCC) and intergenic *f* (TCC).

Given that the TCC*→*T mutation rate is the same in genic and intergenic regions because TCR is ineffective at preventing TCC*→*T mutations in Europeans, we should expect the frequency *f* (TCC) to be slightly higher among private genic SNPs than among private intergenic SNPs. This is because the frequencies of all mutation types sum to 1; mutation types that are efficiently prevented by TCR should have lower frequencies in genic regions than in intergenic regions, and mutation types that are not very susceptible to TCR must have higher genic frequencies to compensate. As predicted by this logic, *f* (TCC) is higher among private genic European SNPs than among private intergenic European SNPs (Figure 3F). In contrast, when PAs and PAf are partitioned into genic and intergenic SNPs, the genic SNP sets have lower TCC*→*T frequencies, suggesting that TCR of this mutation type is relatively efficient in non Europeans. This TCR differential alone could modestly elevate the European TCC*→*T mutation rate. However, it is not likely to be the sole cause of the observed TCC*→*T rate acceleration because this acceleration is evident in both genic and intergenic regions.

## Discussion

It is beyond the scope of this article to pinpoint why the rate of TCC*→*T increased in Europe. However, some promising clues can be found in the literature on ultraviolet-induced mutagenesis. In the mid-1990s, Drobetsky, et al. and Marionnet, et al. each observed that TCC*→*T dominated the mutational spectra of single genes isolated from UV-irradiated cell cultures [29, 30]. Much more recently, Alexandrov, *et al.* systematically inferred “mutational signatures” from 7,042 different cancers and found that melanoma has a unique mutational signature not present in any other cancer type they studied [19]. Melanoma somatic mutations consist almost entirely of C*→*T transitions, 28% of which are TCC*→*T mutations [19, 31]. The mutation types CCC*→*CTC and TCT*→*TTT, two other candidates for rate acceleration in Europe, are also prominent in the spectrum of melanoma (Supporting Information Section S11). Incidentally, melanoma is not only associated with UV light exposure, but also with European ancestry, occurring at very low rates in Africans, African Americans, and even light-skinned Asians [32, 33, 34]. A study of the California Cancer Registry found that the annual age-adjusted incidence of melanoma cases per 100,000 people was 0.8-0.9 for Asians, 0.7-1.0 for African Americans, and 11.3–17.2 for Caucasians [35]. Melanoma incidence in admixed Hispanics is strongly correlated with European ancestry [35, 33, 34].

The association of TCC*→*T mutations with UV exposure is not well understood, but two factors appear to be in play: 1) the propensity of UV to cross-link the TC into a base dimer lesion and 2) poorer repair efficacy at TCC than at other motifs where UV lesions can form [36, 37]. Drobetsky, et al. compared the incidence of UV lesions to the incidence of mutations in irradiated cells and found that TCC motifs were not hotspots for lesion formation, but instead were disproportionately likely to have lesions develop into mutations rather than undergoing error-free repair [29].

Despite the strong evidence that UV causes TCC*→*T mutations, the question remains how UV could affect germline cells that are generally shielded from solar radiation. Although the testes contain germline tissue that lies close to the skin with minimal shielding, to my knowledge it has not been tested whether UV penetrates this tissue effectively enough to induce spermatic mutations. Another possibility is that UV might indirectly cause germline mutations by degrading folate, a DNA synthesis cofactor that is transmitted through the bloodstream and required during cell division [38, 3, 4, 39]. Folate deficiency is known to cause DNA damage including uracil misincorporation and double-strand breaks, leading in some cases to birth defects and reduced male fertility [40, 41, 42]. It is therefore possible that folate depletion could cause some of the mutations observed in UV-irradiated cells, and that these same mutations might appear in the germline of a light-skinned individual rendered folate-deficient by sun exposure. It has also been hypothesized that, in a variety of species, differences in metabolic rate can drive latitudinal gradients in the rate of molecular evolution [43, 44, 45].

Although the data presented here do not reveal a clear mechanism, they leave little doubt that the European population experienced a recent mutation rate increase. TCC*→*T and a few other C*→*T transitions exhibit the clearest evidence of European rate acceleration, but other mutation types might have experienced smaller rate changes within Europeans or other human populations. Pinpointing finer-scale mutation rate changes will be an important avenue for future work.

Even if the overall European mutation rate increase was small, it adds to a growing body of evidence that molecular clock assumptions break down on a faster timescale than generally assumed during population genetic analysis. It was once assumed that the human lineage’s mutation rate had changed little since we shared a common ancestor with chimpanzees, but this assumption is losing credibility due to the conflict between direct mutation rate estimates and molecular-clock-based estimates [8, 9]. Although this conflict might have arisen from a gradual decrease in the rate of germline mitoses per year as our ancestors evolved longer generation times [12, 13], the results of this paper indicate that another force may have come into play: change in the mutation rate per mitosis. If the mutagenic spectrum was able to change during the last 60,000 years of human history, it might have changed numerous times during great ape evolution and beforehand. Given such a general challenge to the molecular clock assumption, it may be wise to infer demographic history from mutations such as CpG transitions that accumulate in a more clocklike way than other mutations [8, 20]. At the same time, less clocklike mutations may provide valuable insights into the changing biology of genome integrity.

## Methods

Publicly available VCF files containing the 1000 Genomes Phase I data were downloaded from www.1000genomes.org/data. Ancestral states were inferred using the UCSC alignment of the chimp PanTro4 to the human reference genome hg19. These data were then subsampled to obtain four sets of SNPs: PE (derived allele private to Europe), PAs (derived allele private to Asia), PAf (derived allele private to Africa), and PAsE (fixed ancestral in Africa but variable in both Asia and Europe).

### 1 Construction of private SNP sets PE, PAs, PAf, and PAsE

The definitions of PE, PAs, and PAf differ slightly from the definitions of continent-private SNPs in the manuscript announcing the release of the 1000 Genomes Phase I data [18]. In that paper, a SNP is considered private to Africa if it is variable in at least one of the populations LWK (Luhya from Kenya), YRI (Yoruba from Nigeria), and ASW (African Americans from the Southwestern USA). In contrast, I consider a SNP to be private to Africa if it is variable in either LWK or YRI and is not variable in any of the following samples: CHB (Chinese from Beijing), CHS (Chinese from Shanghai), JPT (Japanese from Tokyo), CEU (Individuals of Central European descent from Utah), GBR (Great Britain), IBS (Spanish from the Iberian Peninsula), TSI (Italians from Tuscany), and FIN (Finnish). A private African SNP might or might not be variable in any of the admixed samples ASW, MXL (Mexicans from Los Angeles), CLM (Colombians from Medellin), and PUR (Puerto Ricans). Similarly, a private European SNP in PE is variable in one or more of the CEU, GBR, IBS, TSI, and FIN, is not variable in any of YRI, LWK, CHB, CHS, or JPT, and might or might not be variable in ASW, MXL, CLM, and PUR. The private Asian SNPs in PAs are variable in one or more of CHB, CHS, or JPT, are not variable in any of YRI, LWK, CEU, GBR, IBS, TSI, and FIN, and might or might not be variable in ASW, MXL, CLM, and PUR. These definitions are meant to select for mutations that have been confined to a single continent for most of their history except for possible recent transmission to the Americas. The shared European-Asian SNPs in PAsE are variable in one or more of CHB, CHS, or JPT plus one or more of CEU, GBR, IBS, TSI, and FIN and are not variable in YRI or LWK. Singletons are excluded to minimize the impact of possible sequencing error, and variants with imputation quality lower than RSQ = 0.95 are excluded to minimize erroneous designation of shared SNPs as population-private.

### 2 Statistical analysis of frequency differences

Given two SNP sets *P*_1_ and *P*_2_ and one SNP type *m*, a Pearson’s *χ*^2^ test was used to measure the significance of the difference between the frequency of *m* in *P*_1_ and the frequency of *m* in *P*_2_.

Let *C*_*Pi*_ (*m*) denote the number of type-*m* SNPs in set *P*_*i*_, and let 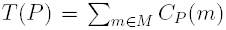 denote the total number of SNPs in *P*. The expected values of *C*_*P1*_ (*m*) and *C*_*P2*_ (*m*), assuming no frequency differences between *P*_1_ and *P*_2_, are calculated as follows based on the 4 *×* 4 contingency tables in Figure 1C,D:

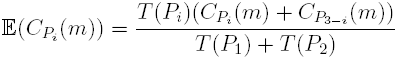

The following *χ*^2^ value with one degree of freedom measures the significance of the difference between *f*_*m*_(*P*_1_) and *f*_*m*_(*P*_2_):

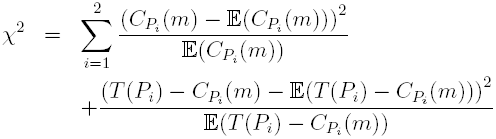

### 3 Nonparametric bootstrapping within chromosomes

To assess the variance of *f* (TCC) within each of the autosomes and the X chromosome, each private SNP set PE, PAs, and PAf was partitioned into non-overlapping bins of 1,000 consecutive SNPs. The frequency *f* (TCC) of the mutation TCC*→*T was computed for each bin and the distribution of these estimates is shown in Figure 1C. No partitioning into separate bins was performed for chromosome Y because the entire chromosome has only 1,130 private European SNPs, 1,857 private Asian SNPs and 3,852 private African SNPs. Instead the global frequency of TCC*→*T was computed for each SNP set restricted to the Y chromosome.

### 4 Quantifying strand bias

Gene locations and transcription directions for hg19 were downloaded from the UCSC Genome browser. For the purpose of this analysis, each SNP located between the start codon and stop codon of an annotated gene is considered to be a genic SNP. All SNPs not located within introns or exons are considered to be intergenic SNPs.

Within each private SNP set P, each mutation type *m* with an A or C ancestral allele was counted separately on transcribed and non-transcribed genic strands to obtain counts **T**(P*, m*) and **N**(P*, m*). (Each mutation with a G/T ancestral allele on the transcribed strand is equivalent to a complementary A/C ancestral mutation on the non-transcribed strand.) The strand bias of mutation *m* is then defined to be **S**(P*, m*) = **N**(P*, m*)*/***T**(P*, m*). The significances of the strand bias differences **S**(PAf*, m*) *-* **S**(PE*, m*) and **S**(PAf*, m*) *-* **S**(PAs*, m*) were assessed using a *χ*^2^ test with 1 degree of freedom. Assuming no difference in strand bias between *P*_1_ and *P*_2_, the expected numbers of transcribed-strand and nontranscribed-strand mutations are the following:

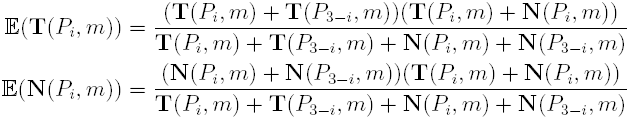

The *χ*^2^ value measuring the significance of the difference between **N**(*P*_1_*, m*)*/***T**(*P*_1_*, m*) and **N**(*P*_2_*, m*)*/***T**(*P*_2_*, m*) is computed as follows:

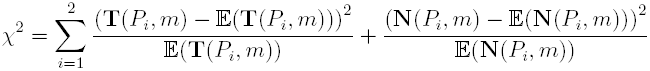

Non-parametric bootstrapping was used to estimate the variance of TCC*→*T strand bias within each population. The transcribed portion of the genome was partitioned into 100 bins containing approximately equal numbers of SNPs, and 100 replicates were generated each by sampling 100 bins with replacement. For each replicate, the frequency of TCC*→*T was calculated on the transcribed and non-transcribed strands. These two frequencies were added together to obtain the cumulative TCC*→*T frequency within genic regions. The distribution of **S**(TCC *→* T) across replicates is shown for each population in Figure 3C.

Bootstrapping was similarly applied to intergenic SNPs by partitioning the non-genic portion of the genome into 100 bins with similar SNP counts. 100 bootstrap replicates were generated by sampling 100 bins with replacement, and the intergenic TCC*→*T frequency was computed for each replicate.

The distribution of ratios in Figure 3D was generated by pairing up each genic bootstrap replicate with a unique intergenic bootstrap replicate and calculating the ratio of genic *f* (TCC) to intergenic *f* (TCC), thereby obtaining 100 estimates of the ratio *f*_genic_(TCC)*/f*_intergenic_(TCC).

## Acknowledgements

I am grateful to Rasmus Nielsen for advice and manuscript comments, and to two anonymous reviewers for providing feedback that improved upon an earlier draft. David Reich shared valuable insights into gene flow between early European farmers and hunter gatherers, and Richard Durbin, Stuart Linn, and Elad Ziv contributed additional helpful comments and suggestions. This work was supported by a National Science Foundation Graduate Research Fellowship (awarded to K.H.) and NIH grant IR01GM109454-01 (awarded to Rasmus Nielsen, Yun Song, and Steve Evans).

